# Bootstrat: Population Informed Bootstrapping for Rare Variant Tests

**DOI:** 10.1101/068999

**Authors:** Hailiang Huang, Gina M. Peloso, Daniel Howrigan, Barbara Rakitsch, Carl Johann Simon-Gabriel, Jacqueline I. Goldstein, Mark J. Daly, Karsten Borgwardt, Benjamin M. Neale

## Abstract

Recent advances in genotyping and sequencing technologies have made detecting rare variants in large cohorts possible. Various analytic methods for associating disease to rare variants have been proposed, including burden tests, C-alpha and SKAT. Most of these methods, however, assume that samples come from a homogeneous population, which is not realistic for analyses of large samples. Not correcting for population stratification causes inflated *p*-values and false-positive associations. Here we propose a population-informed bootstrap resampling method that controls for population stratification (Bootstrat) in rare variant tests. In essence, the Bootstrat procedure uses genetic distance to create a phenotype probability for each sample. We show that this empirical approach can effectively correct for population stratification while maintaining statistical power comparable to established methods of controlling for population stratification. The Bootstrat scheme can be easily applied to existing rare variant testing methods with reasonable computational complexity.

**Author Summary:** Recent technology advances have enabled large-scale analysis of rare variants, but properly testing rare variants remains a significant challenge as most rare variant testing methods assume a sample of homogenous ethnicity, an assumption often not true for large cohorts. Failure to account for this heterogeneity increases the type I error rate. Here we propose a bootstrap scheme applicable to most existing rare variant testing methods to control for population heterogeneity. This scheme uses a randomization layer to establish a null distribution of the test statistics while preserving the sample genetic relationships. The null distribution is then used to calculate an empirical *p*-value that accounts for population heterogeneity. We demonstrate how this scheme successfully controls the type I error rate without loss of statistical power.

## Introduction

Case-control association studies have successfully identified significant genetic associations across a wide range of phenotypes including inflammatory bowel diseases [1-3], cardiovascular diseases [4-7], and psychiatric diseases [8-11]. For complex traits, the vast majority of significant associations have been largely restricted to common variants (minor allele frequency > 1%). In most instances, identification of causal variants is challenging because of pervasive linkage disequilibrium among common variants in a region. Analysis of rare variants has the potential to nominate additional regions of interest; in particular coding variation holds promise for identifying specific genes of interest. The advent of exome chip and advances in whole exome and genome sequencing allow for high throughput detection and genotyping of rare variants. Finding rare variants underlying human disorders, however, presents a significant challenge due to the limited statistical power when only a small number of individuals carry a given variant.

Several approaches have been designed to improve statistical power through variant aggregation, patterns of observed variation, and mixture models [12-15]; however no approach is inherently designed to control for the confounding of population stratification. Stratification is not an issue when datasets come solely from a single homogenous population, but can become severe as dataset sizes increase and are combined across multiple cohorts. For example, a recent study collected roughly 35,000 European samples across multiple cohorts and had to use 15 principal components (PCs) to correct for population stratification [2,16]. Furthermore, stratification of rare variants may operate on a finer population scale than is easily correctable using current methods [17]. Not appropriately adjusting for stratification can lead to inflated test statistics and false-positive associations, limiting the usage of these methods.

Population stratification in common variant association tests can be controlled by using genetic principal component analysis (PCA) [8] or linear mixed models (LMM) [18,19]. PCA models ancestry differences among individuals using principal components of thousands of genetic markers across the genome, which are then used as covariates in association tests. LMM uses the random effect model to estimate the contribution from the genetic relationship matrix (GRM) to phenotypic variance, and adjusts the association test statistics accordingly. While both methods work well for common variant association tests, extending them to rare variants, if possible, is not obvious because of their complexity.

Several existing methods have the potential to be adapted to control the population stratification in rare variant association tests but they each have their pitfalls. The genomic control method [20] adjusts the association test statistics using the inflation factor estimated from unlinked variants. Despite its simplicity, the assumption that the inflation factor is constant across the genome is unlikely to be true especially for rare variants, therefore limiting the use of this method. Pritchard et al. proposed a model-based clustering approach to infer the population structure [21] and took advantage of this structure to perform randomizations for calculating the significance of associations [22,23]. This method can be retrofit to existing rare variant tests by clustering individuals of close genetic distance into a subgroup and shuffling phenotypes uniformly within each group (discussed in **Results** and **Methods**). However, this approach is not optimal because genetic relatedness falls along a broad continuum and can be hard to categorize into a finite number of subgroups. In addition, some clusters may only have cases or controls and thus have to be dropped, resulting in a smaller effective sample size. A more sophisticated approach [24] used the Fisher’s non-central hypergeometric distribution [25] to resample disease status such that the odds of a subject being selected as a case are equal to his or her odds of disease conditional on confounder variables. Because disease status was used to train the model, a potential problem is that the empirical distribution of test statistics can be biased, leading to a biased empirical *p*-value (discussed in **Results**).

We propose an alternative approach, Bootstrat, that addresses the limitations of the aforementioned methods using a bootstrap based randomization scheme informed by the magnitude of genetic distance learned from the dataset. This scheme assigns higher probabilities of sampling phenotypes from genetically related individuals. Unlike permutation within discrete clusters, we are able to include all samples in the randomization procedure. The randomization process is driven only by the genetic similarity and has the potential to deal with non-monotonic effects across ancestry space. When applied to a simulated dataset, Bootstrat successfully controls the genomic inflation due to population stratification in both single variant and gene-based tests, while keeping similar computation efficiency. In gene-based tests, Bootstrat shows improved type I error control over other methods while maintaining the same statistical power.

## Results

### Dataset

All analyses in this study were performed on a dataset of 2,000 samples. Half of the samples (1,000) were randomly selected from an American control cohort and the other half were randomly selected from a Swedish control cohort (**Methods::Dataset**). To simulate the effect of population structure, we randomly selected 70% of American samples and 30% of Swedish samples as cases. The remaining samples were labeled as controls. For gene-based analyses, we selected variants with a minor allele frequency <5% and labeled as non-synonymous, splice-site, or nonsense. Genes with at least 40 minor alleles were used for assessment. This threshold was chosen to allow good asymptotic behavior and enough statistical power [26]. There were 5,725 genes and 44,392 variants available for analysis. The number of minor alleles per gene ranges from 40 to 3,506 with mean of 146 and median of 105.

### Bootstrat

Typically, to empirically evaluate the significance of a hypothesis, a permutation test can be used to construct a null distribution of the test statistics. The permutation test is non-parametric (i.e. distribution free) and can provide an exact significance level. However, it has an important assumption that all samples are exchangeable under the null hypothesis [27], i.e., the joint probability distribution of the permuted sequence is the same as that of the original sequence. This assumption holds in a homogenous population where samples have equal probabilities to permute with each other. In a stratified dataset, this assumption no longer holds because permutations between samples of the same ethnicity have higher exchange probabilities than those between samples of different ethnicities. For example, in a dataset with mostly European samples, if A has a nontrivial amount of Asian ancestry composition and B is almost 100% European, the probability for A to exchange with B would be higher than that for B to exchange with A. This is because sample B is more likely to exchange with a sample of his/her own ancestry but sample A can only exchange with samples in other ancestries. Therefore, the probability for sample A to shuffle with sample B can be different from the probability for sample B to shuffle with sample A.

When this happens, bootstrap can be used in place of the permutation test to evaluate the significance of a hypothesis. Weighted bootstrap [1-3,28] relaxes the exchangeability condition [1,3-7] by weighting each sample in the original dataset and resampling them randomly with replacement. Therefore, we use weighted bootstrap, instead of the permutation with uniform probabilities, to resample phenotypes and create a null distribution of the association test statistics with awareness to population stratification. The weighting reflects the genetic distances between samples, such that a sample is more likely to draw a phenotype from samples sharing the similar ethnicity. This is intuitively illustrated in Fig 1, which shows two samples shuffle only with other samples that are genetically close. Once the null distribution corrected for population stratifications is established, it can be used to calculate the empirical significance of the observed test statistic.

**Fig 1.**
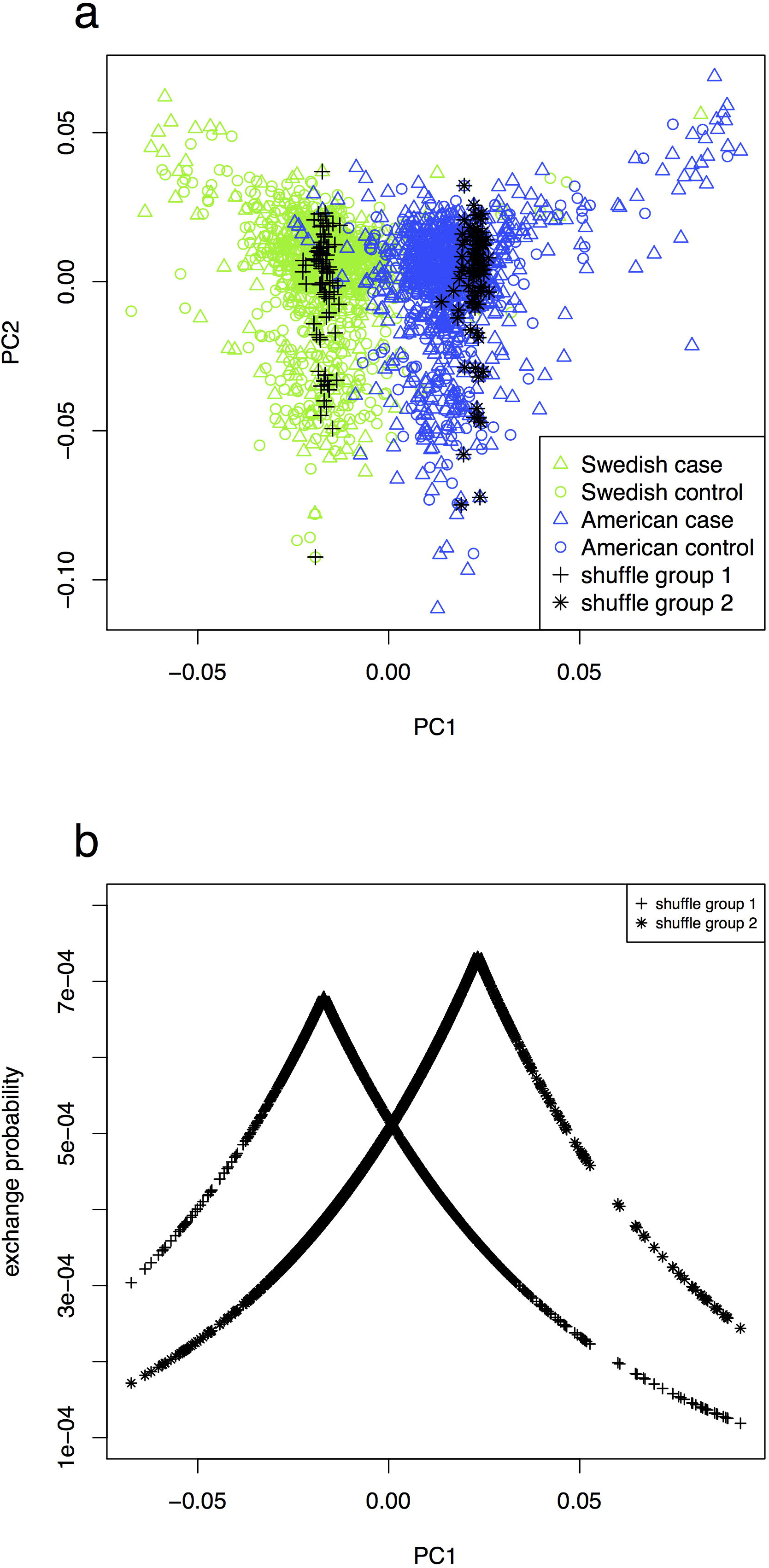
Population stratification was simulated using Swedish and American control cohort. (a) Case and control statuses were simulated for Swedish and American cohorts (**Methods**). The first two principal components were plotted, although PC1 alone was sufficient to control the population stratification (**Results::Principal components**). We traced 100 randomizations of two randomly picked samples (as back crosses and asterisks respectively) as an illustration, showing that samples are shuffled only with other samples that have close genetic distance. (b) The exchange probability for the two randomly picked samples decreases with the genetic distance.

### Measure of population stratification

In a GWAS with no confounding effects (e.g. no population stratification), most variants are not associated with the trait and their test statistics follow the null distribution. Presence of confounding effects inflates the test statistics in the study and causes them to deviate from the null distribution. The Quantile-Quantile (QQ) plot compares the observed quantiles versus the expected quantiles under the null hypothesis and is frequently used to check that the distribution of statistics follows expectation. For common variants, a well-behaved QQ plot only departs from the diagonal line at the tail, representing the small proportion of significant associations. In addition to the QQ plot, the genomic inflation factor, usually denoted as λ, quantitatively measures the deviation of the observed distribution from the null distribution. λ is defined as the ratio of the median test statistic from the study over the median test statistic from the null distribution and is expected to be 1 for common variants when there is no population stratification. Both QQ plot and the genomic control factor work well for common variants in measuring population stratification, but are not optimal for rare variants, which usually have different asymptotic behaviors and will be discussed in later sections. Unless otherwise noted, the test statistic used in this study is from the single degree of freedom chi-square test using single variant logistic regression.

### Principal components

The first principal component (PC) was able to identify the two population cohorts in our dataset (Fig 1), and controls the population stratification in the QQ plot (S1 Fig) with a reduced genomic inflation factor (λ) from 1.330 to 1.000. Increasing the number of PCs to 2 and 3 has λ=1.010 and 0.992, respectively. Thus, using more than the first PC does not improve our control over population stratification effects, and hurts statistical power [4,8-11]. We therefore use only the first PC throughout this study.

### Estimate the Bootstrat parameter

Bootstrat has a parameter, *γ*, that controls the magnitude of the correction. It resembles the grid size or the number of clusters in the “permutation within cluster” approach (See **Methods**). A larger *γ* results in a stronger control of stratification and should be used in a severely stratified dataset. We estimate the optimal *γ* by doing a grid search to find the smallest *γ* that sufficiently controls the population stratification (S2 Fig). We note that further increasing *γ* beyond the optimal value resembles further dividing population homogenous groups into smaller subgroups, which does not achieve better population control and results in lower power. The desired level of genomic control can be variable depending on many factors such as sample size and genotyping chip. We took the genomic control factor from common variant association tests using PC as covariates as the desired level. In the grid search, we allowed λ to fall within the 1% range of this desired level. In this study, the desired level of λ is 1.000 (Table 1) and we allow λ to range between 1.010 and 0.990. We calculated λ for *γ* between -3 and 8 using common variants (Table 1). We found Bootstrat using *γ* =4 reported λ of 1.002, indicating a perfect control of inflation (from λ=1.323 in the ordinary permutation). Visual inspection of the QQ plot also confirms the full control of stratification for common variants (S2 Fig).

**Table 1.**
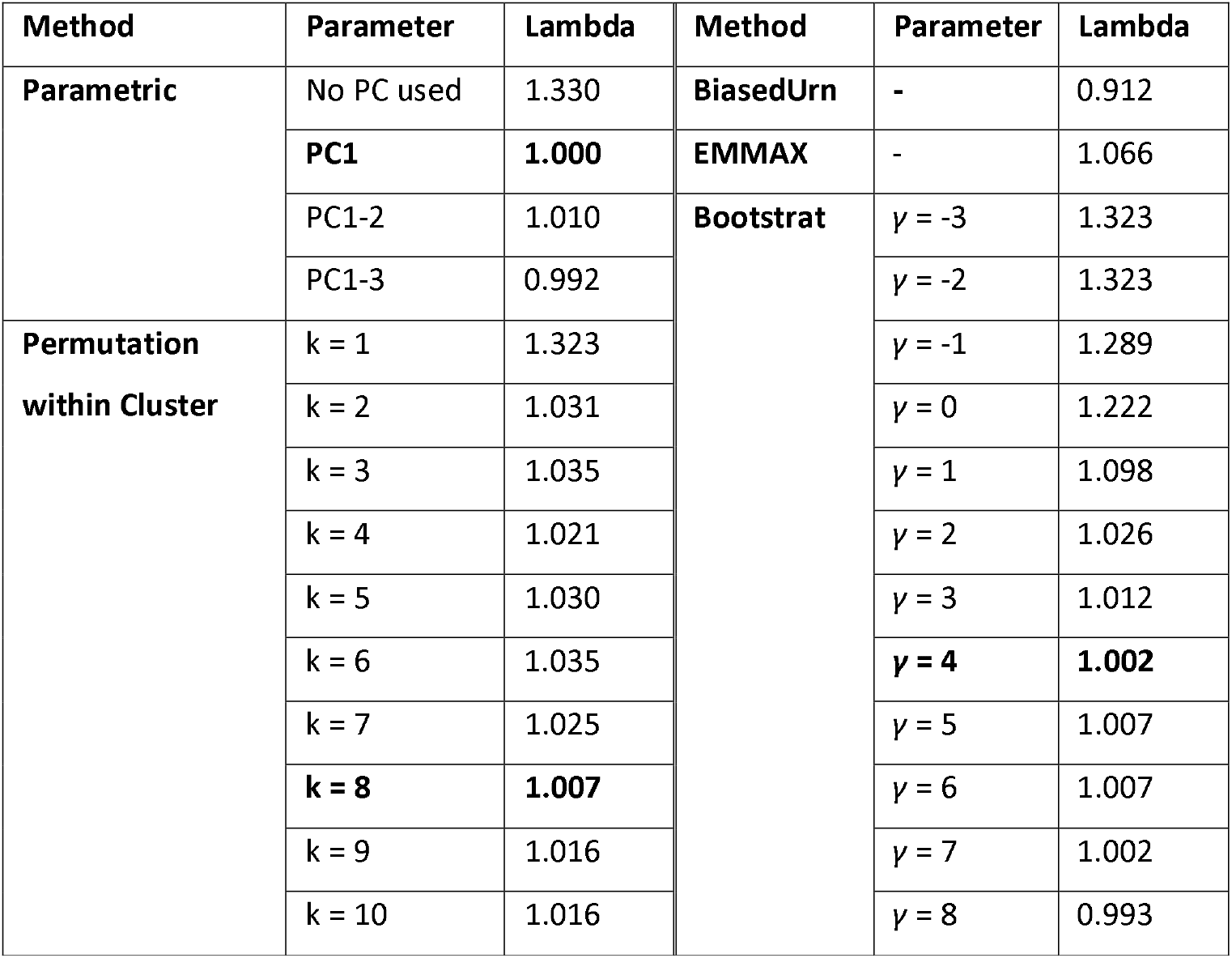
Genomic inflation factor using common variants. k is the number of clusters used in the permutation within cluster approach. *γ* is the parameter used in Bootstrat (**Methods::Bootstrat**).

### Estimate the number of clusters

A naïve approach to randomize population-stratified samples is to group them into ethnicity clusters and permute uniformly within each cluster [21-23]. The number of clusters can be estimated similarly as estimating *γ* for Bootstrat: we chose the smallest number of clusters that sufficiently controls the stratification. Using the same desired genomic control level (λ ranges between 1.010 and 0.990), we found that dividing the samples into 8 clusters achieves the optimal genomic control (Table 1).

### Single common variant test

Common variants were defined as variants with minor allele frequency (MAF) > 5%.22,415 common variants are available in this study after QC. Parametric test adjusting for PCs of ancestry is the *de facto* method for controlling the population stratification, and has been demonstrated to work well [8,10]. We show that Bootstrat is able to control the population stratification as good as the parametric test with PC adjustment (Fig 2). Genomic inflation factors are 1.000 for parametric test with PCs, 1.002 for Bootstrat with *γ*=4, 1.007 for permutation within cluster (8 clusters), 0.912 for BiasedUrn and 1.066 for EMMAX. Note that the purpose of this section is not to find the best method in controlling stratification in single common variant tests. We consider this a solved problem: parametric tests using PCs or LMM[18,19] have been proven to be the solutions. We instead used this established solution as gold standard to estimate parameters in other approaches, including Bootstrat and the permutation within cluster approach. It is therefore not fair to use the single common variant test as the metric to perform method comparison.

**Fig 2.**
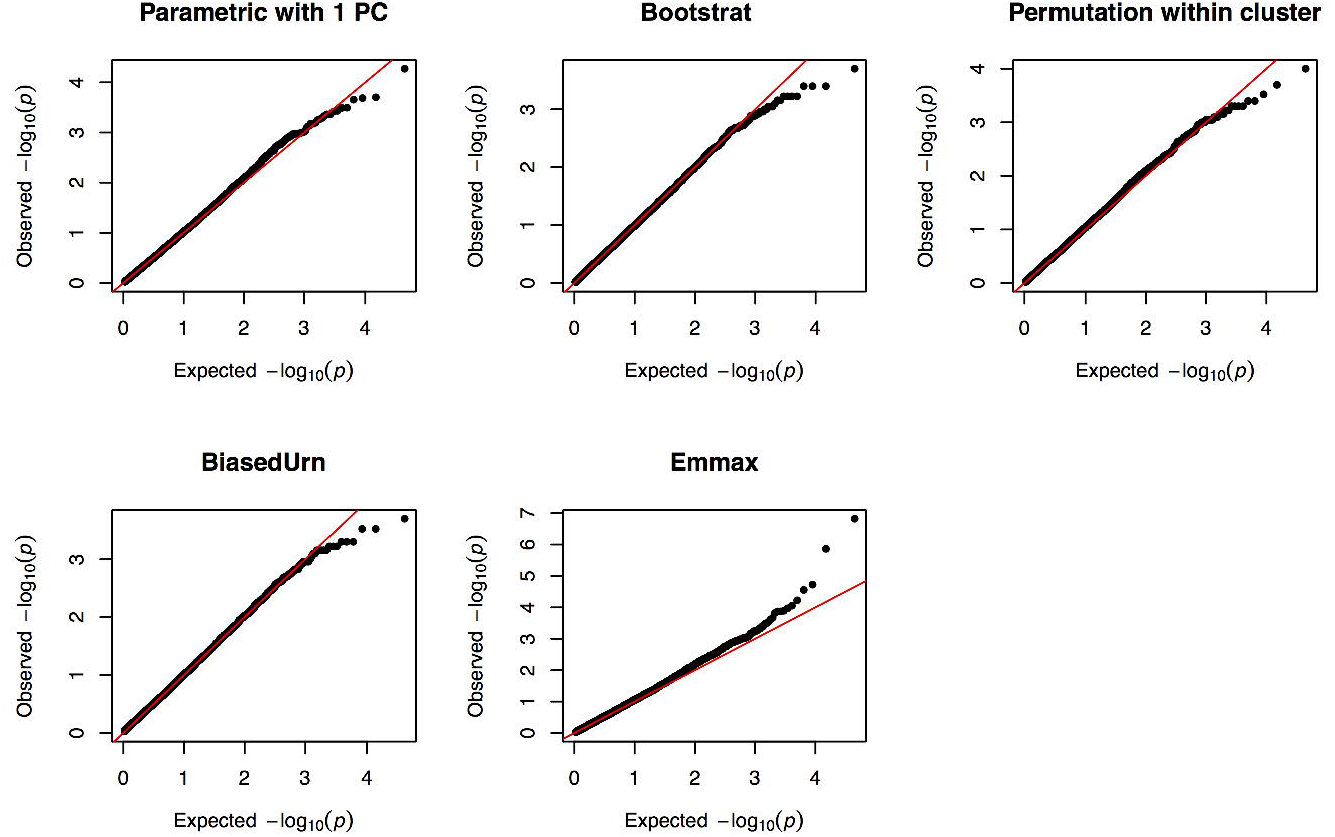
QQ plot of common variants. QQ plot of the test statistics of common variants for various methods discussed. For methods with parameters, we used the parameter that achieves the best performance (**Results**): we used *γ*=4 for Bootstrat and 8 clusters for permutation within cluster.

### Single rare variant test

Rare variants were defined as variants with MAF < 1% but with at least 10 minor alleles (MAF = 0.25% in the current sample). There were 9,881 rare variants available in this study after QC. Under the case/control study design, the test statistic distribution among rare variants often fails to meet asymptotic assumptions of chi-square distributed tests [26]. As a result, the corresponding p-value distribution lacks uniformity, making visual inspection of the QQ plot uninformative and introducing fundamental bias to measures of λ. To address this problem, we generated an empirical null distribution of test statistics via permutation (see **Methods:: Empirical null distribution for single rare variant tests**). These permutations are drawn independent from population structure and have the advantage of conditioning on the sample size, selected allele frequency distribution and case-control balance within the dataset. Both the QQ plot and λ can be recalibrated using this permuted null distribution.

Overall, every method reduces the genomic inflation present among rare variants confounded by population stratification, whereby no control for population stratification leads to a comparable level of genomic inflation as for common variants (λ=1.356). Fig 3 shows the empirical QQ plot and corresponding median λ statistic for single rare variants under the variety of methods analyzed. Visual inspection of the QQ plot against the asymptotic chi-square shows how the BiasedUrn test is generally deflated across the distribution of test statistics, while the parametric test controlling for the 1st PC shows marked deflation at the tail end of the distribution, suggesting a lack of power due to the inclusion of a covariate into the regression. After recalibrating observed test statistics to their permuted null distribution, tests generally track more closely to expectation across the board. While no test fully accounts for population stratification among rare variants, Bootstrat outperforms all other tests when measuring genomic inflation, matching the parametric test controlling for the 1st PC under the asymptotic test distribution (λ=1.047), and showing the lowest inflation using the empirical test distribution (λ=1.037).

**Fig 3.**
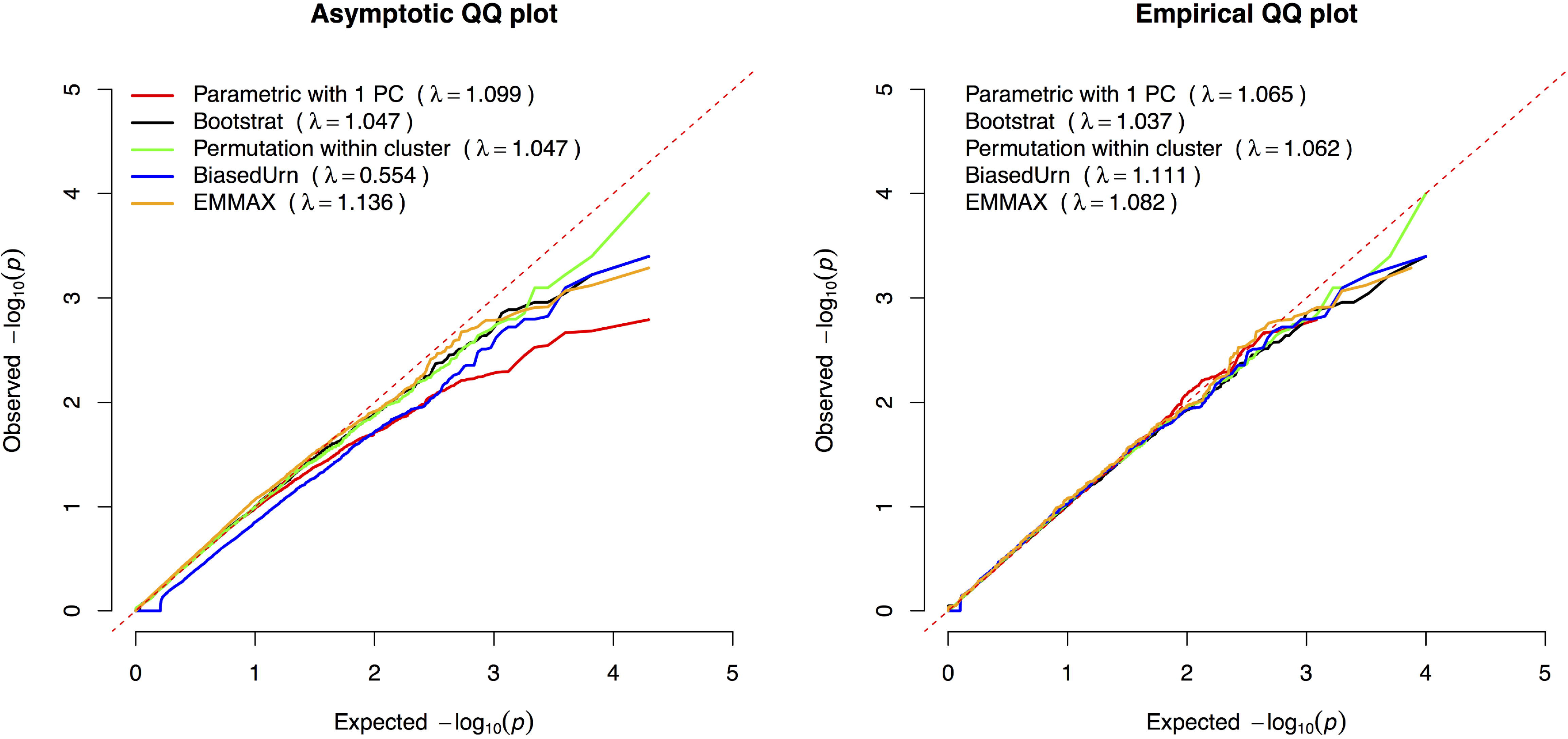
QQ plot of rare variants. QQ plot for various methods calibrated using the null phenotype. Only rare variants were plotted. We used the same parameters as for common variants: *γ*=4 for Bootstrat and 8 clusters for permutation within cluster.

### Gene-based rare variant test

Variant-set Kernel Association Test (SKAT) [2,12,16] uses a kernel machine to test for the association between a set of SNPs within a gene and an outcome. The set of variants that are tested together is defined based on variant characteristics such as frequency and *in silico* functional class. SKAT using permutations for assessing significance was modified to incorporate the ancestry informative randomization from Bootstrat. There were 5,725 genes tested with an outcome simulated to have population structure using 5 models: (1). SKAT with empirical significance, (2). SKAT with random permutation for significance, (3). SKAT adjusted for PCs of ancestry as covariates, (4). SKAT adjusted for PCs of ancestry as covariates with random permutation, and (5). SKAT modified with Bootstrat. Inflation in the Q-Q plot was observed when no adjustment for population has been performed (models 1 and 2), and adjusting for population structure by using PCs of ancestry or Bootstrat more appropriately controls for the population structure (Fig 4).

**Fig 4.**
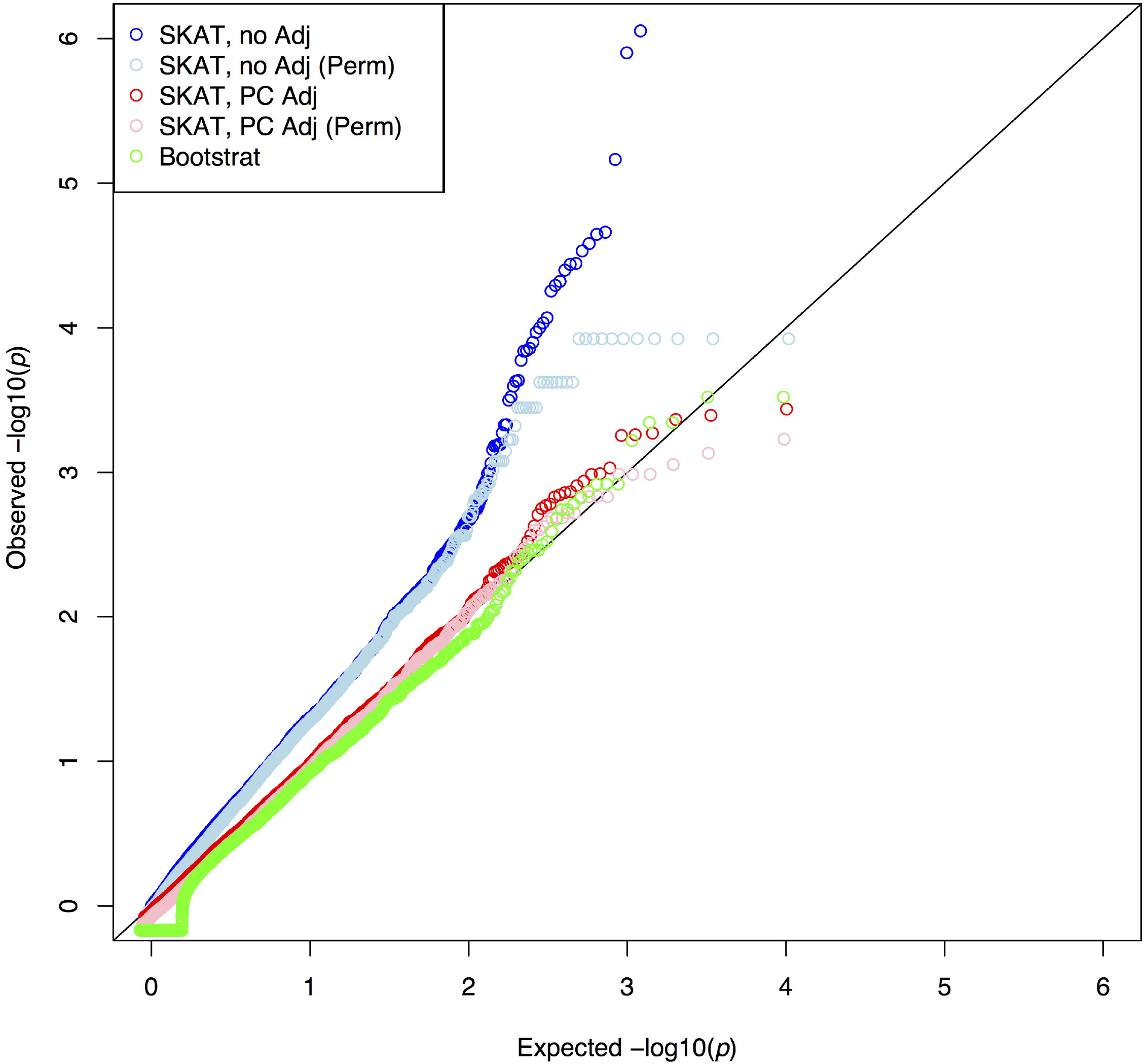
QQ plot of gene-based tests under the null. QQ plot of the test statistics of gene-based tests. “SKAT, no Adj”: SKAT with empirical significance. “SKAT, no Adj (Perm)”: SKAT with random permutation for significance. “SKAT, PC Adj”: SKAT adjusting for PC1 as a covariate. “SKAT, PC Adj (Perm)”: SKAT adjusting for PC1 as a covariate with random permutation for significance. “Bootstrat”: SKAT modified with Bootstrat.

We assessed the type I error and power for the gene-based test using the 5 models described above. We found that type I error rates were severely inflated when we did not account for the population structure (models 1 and 2). The inflation was not fully reduced when PCs of ancestry were used for adjustment of population structure (Fig 5). In contrast, Bootstrat was able to better control the type I error at the significance levels of 0.05, 0.01 and 0.005. For significance levels of 0.001 and 0.0005, Bootstrat has similar performance as using PCs. Furthermore, power was found to be similar across all the models at various nominal type I errors (Table 2). Note that although Bootstrat has an additional parameter, this parameter was trained using single common variant test and stayed the same across rare variant tests, including the single variant test and the gene-based test. Therefore, we argue that the superior performance of Bootstrat in testing rare variants is not due to overfitting.

**Fig 5.**
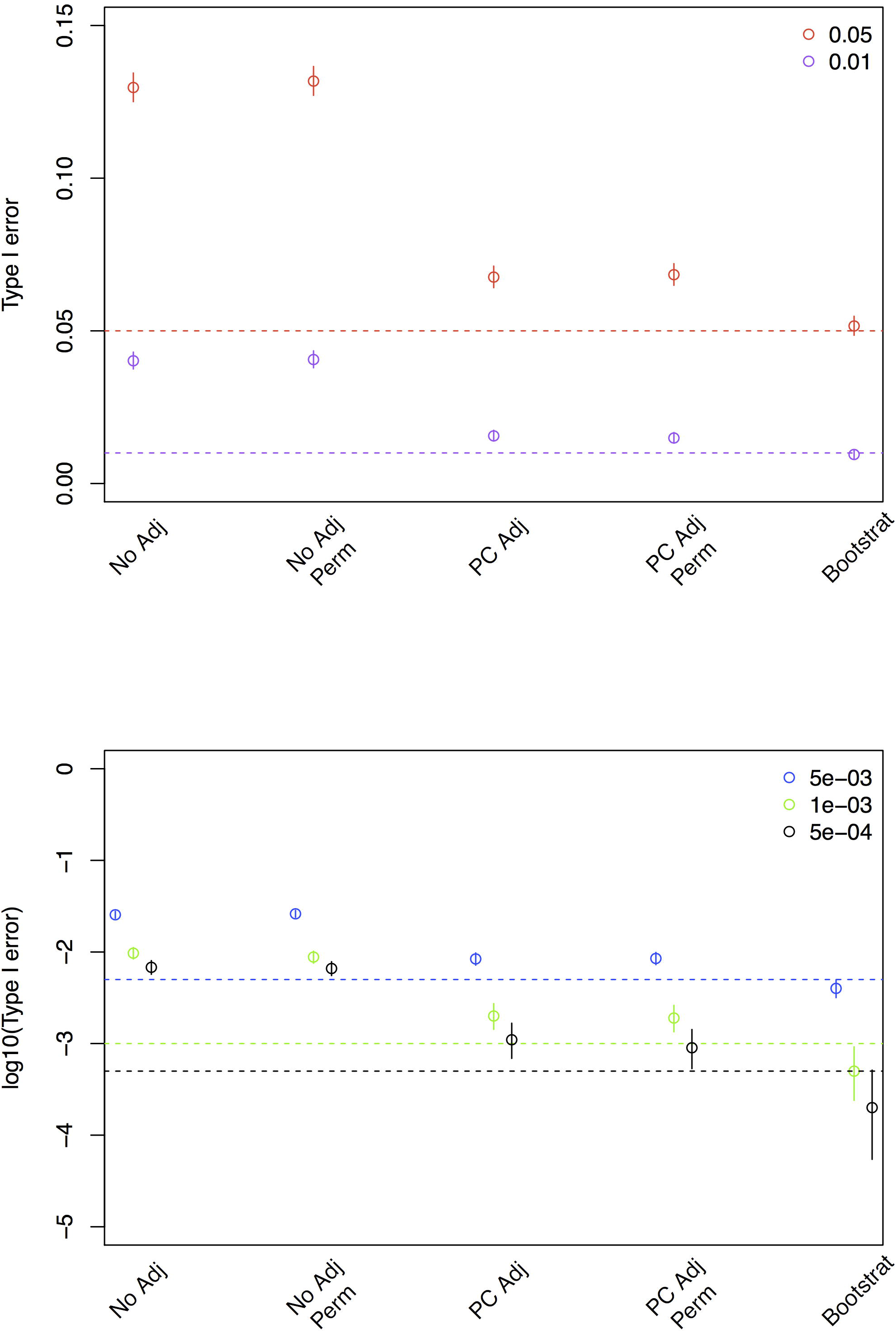
Type I error of gene-based tests using rare variants. Type I error was calculated as the proportion of genes meeting the specified α level (color coded) across all the genes analyzed. Dashed lines indicate the expected type I error and open circles indicate the observed type I error for each method.

**Table 2.** Power for gene-based tests using rare variants. Power was assessed using an outcome simulated with population structure and an odds ratio of 2.5 and 5 (**Methods::Type I Error and Power for and gene-based tests**). Prop. functional is the proportion of variants that are functionally related to the outcome. We assigned each variant to be neutral or have an effect with a 30% (or 50%) probability. A greater proportion of functional variants simulates less noise in the gene-based association testing. α is the type I error at which the power was assessed. Power and its standard error were both reported.

| OR | Prop. func. | $\alpha$ | No adj | No adj (perm) | PC adj | PC adj (perm) | Boostrat |
| --- | --- | --- | --- | --- | --- | --- | --- |
| 2.5 | 0.3 | 0.05 | 0.51 ± 0.03 | 0.52 ± 0.03 | 0.45 ± 0.03 | 0.45 ± 0.03 | 0.42±0.03 |
| 2.5 | 0.3 | 0.01 | 0.36 ± 0.03 | 0.37 ± 0.03 | 0.34 ± 0.03 | 0.34 ± 0.03 | 0.30±0.03 |
| 2.5 | 0.5 | 0.05 | 0.66 ± 0.03 | 0.66 ± 0.03 | 0.65 ± 0.03 | 0.65 ± 0.03 | 0.58±0.03 |
| 2.5 | 0.5 | 0.01 | 0.54 ± 0.03 | 0.53 ± 0.03 | 0.52 ± 0.03 | 0.52 ± 0.03 | 0.45±0.03 |
| 5 | 0.3 | 0.05 | 0.70 ± 0.03 | 0.70 ± 0.03 | 0.68 ± 0.03 | 0.68 ± 0.03 | 0.63±0.03 |
| 5 | 0.3 | 0.01 | 0.60 ± 0.03 | 0.60 ± 0.03 | 0.59 ± 0.03 | 0.59 ± 0.03 | 0.53±0.03 |
| 5 | 0.5 | 0.05 | 0.86 ± 0.02 | 0.85 ± 0.02 | 0.86 ± 0.02 | 0.86 ± 0.02 | 0.81±0.02 |
| 5 | 0.5 | 0.01 | 0.77 ± 0.03 | 0.78 ± 0.03 | 0.77 ± 0.03 | 0.77 ± 0.03 | 0.73±0.03 |

### Computational performance

Bootstrat was efficiently implemented: for our study cohort of 2,000 samples, it takes 0.1 minutes to calculate the exchange probability matrix, and 8.8 minutes to perform the permutations using this matrix (Table 3). This time is longer than the ordinary permutation because the probabilistic sampling is more time consuming than sampling with uniform probabilities. However, the time for running randomizations can be reduced significantly (from 8.8 to 3.8 minutes in this study) for case/control study designs using a more efficient design (**Methods:: Efficient implementation for the case-control study design**). The exchange probability matrix only needs to be calculated once per dataset, and can be reused with different phenotypes and parameters. In practice, the run time can be further reduced by using adaptive permutations [16,24] and parallelization platforms such as MapReduce [25,29].

**Table 3.** Run time for each method. CPU time (in minutes) for performing tests on chromosome 1. ‘Preparation time’ is the time for pre-calculating inputs for some methods, including the kinship matrix for EMMAX, clustering for ‘permutation within cluster’, and exchange probability matrix for Bootstrat. This step only needs to be done once per cohort and can be reused in analysis with different phenotypes and options. ‘Analysis time’ is the time for actually performing the association study given the pre-calculations.

| Method | Preparation time | Analysis time | Total |
| --- | --- | --- | --- |
| Parametric | - | 0.4 | 0.4 |
| BiasedUrn | - | 335 | 335 |
| EMMAX | 5.8 | 1.0 | 6.8 |
| Permutation within cluster | 0.0 | 3.3 | 3.3 |
| **Bootstrat** | **0.1** | **8.8** | **8.9** |
| Bootstrat (case/control) | 0.1 | 3.8 | 3.9 |
| SKAT with PC adjustment<br>and permutations | - | 23.0 | 23.0 |
| SKAT with Bootstrat | 0.1 | 33.2 | 33.3 |

## Discussion

We have presented Bootstrat, a bootstrap based randomization scheme that corrects for population stratification in testing rare variants for association with diseases. We compared this scheme with existing methods including EMMAX, BiasedUrn, permutation within cluster and SKAT and found that Bootstrat controls the type I error better than existing methods, while keeping the power at similar levels.

In our comparison, we did not include the recently developed KM method, which is an extension of SKAT to accommodate related subjects from multiple independent families [26,30]. The authors of the KM test only evaluated their method for common variants in SNP sets with families, while we propose to use Bootstrat when there are multiple sub-population groups within the data. Therefore, comparison with KM method is beyond the scope of this paper.

Bootstrat is compatible with both case/control studies and quantitative traits and works with most existing rare variant tests. Users have the option to use this permutation scheme to run any available tests implemented in PLINK. We have implemented this permutation scheme in PLINK, and R code that implements this scheme in the SKAT framework is available upon request.

## Methods

### Dataset

Both cohorts used in this study were population controls genotyped on the exome chip. After QC and removing related samples, the NIMH control cohort has 1,036 samples and 230,329 variants [31]; and the Swedish cohort has 6,063 samples and 246,843 variants [32]. After merging, we kept 199,928 autosomal variants available in both cohorts. We randomly selected 1,000 samples from each cohort to create the dataset used in this analysis (Fig 1).

### Remove related samples

We performed LD prune on the common variants (minor allele frequency > 5%) using PLINK [27,33]. These LD pruned common variants were then used to calculate the Identity-by-descent (IBD) matrix. Using the IBD matrix, we identified 139 related sample pairs (Pi_hat > 0.2) in the Swedish cohort. No related samples were found within the NIMH cohort or across the two cohorts. In the Swedish cohort, we randomly removed one sample in each pair, a total of 134 unique samples, to create a dataset of independent samples.

### Bootstrat

Individuals having the same genetic ethnicity have been shown to cluster closer in the principal components (PCs) space. Many genome-wide association studies (GWAS) used PCs as covariates to correct for population stratification. Various measures have been used to choose which PCs to include in the analyses such as the visual inspection of the QQ plot, the genomic inflation factor, and testing the associations between PCs and case/control. In the following context, we assume users know the appropriate PCs to use from the ordinary GWAS. Denote the p-th PC as *w*_p_ and its eigenvalue as *λ*_p_, we calculated the pairwise Euclidean distance between samples *i* and *j* as

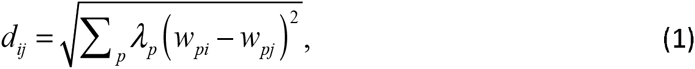

in which the summation of p includes all PCs that one would use in a GWAS. We calculated the raw pairwise exchange probability from this distance using the exponential transformation 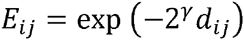 and normalized the raw probability to obtain the final probability for the weighted bootstrap. Other transformations, including linear and inverse, were also considered but we found the exponential transformation provided the most robust results (results not shown). For sample *i*, the probability to draw the phenotype from sample *j* is

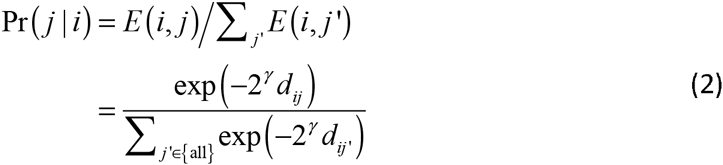

The denominator is the sum of raw pairwise exchange probabilities over all samples in the studies, including sample *i* itself. *γ* is a scaling parameter that controls how the probability scales with the distance, which can take any value between −∞ and +∞. *γ* =-∞ gives uniform exchange probability for each sample pair and is essentially the ordinary bootstrap. *γ* =+∞ confines each sample to only swapping with itself. We performed a grid search to find the optimal *γ* based on the genomic inflation factor using common variants. This is computationally feasible because only the LD pruned common variants (less than 20,000 variants in exome chip) are needed to calculate the inflation factor.

We calculated the test statistics in each bootstrap and created a null distribution for each variant. The empirical *p*-value of an association is defined as the fraction of time a better or equal test statistic was achieved under the null. In practice, we performed bootstrap in an adaptive manner to save computation time [28,33].

### Efficient implementation for the case-control study design

For case-control study design, instead of performing bootstrap for each sample, a more efficient method would be to calculate the probability for a particular sample to be case or control, i.e.,

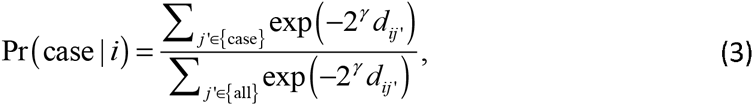

and use this probability to simulate the case/control status from a binomial distribution.

### Baseline methods

As the comparison baseline, we performed logistic regression with and without the PCs as covariates, and permutation tests with a minimum of 1,000 permutations in PLINK 1.08 [1,3,33]. We performed a linear mixed model implementation, EMMAX, using the package released on March 2007 with default parameters [4,19], and Biased Urn using the R ‘BiasedUrn’ package in CRAN with recommended revisions from the paper [8,10,24]. Other methods discussed are:

### Permutation within cluster

This approach divides samples into clusters based on their genetic similarity. In this study, the similarity was calculated as the Euclidian distance using the 1st principal component. Within each cluster, we assume population homogeneity and shuffle samples with uniform probability. Clusters with only case or control samples are ignored. The number of clusters was chosen such that the genomic control has achieved a satisfactory level (**Results::Estimate the number of clusters**).

### SKAT

We used The Sequence Kernel Association Test (SKAT) [12] version 0.81 (implemented in R) to for the gene-based tests using an identity-by-state kernel. SKAT evaluates qualitative and quantitative traits by comparing rare variant similarity with phenotypic similarity, thus permitting variants to be associated with an increase and decrease in the phenotype within a gene.

### Empirical null distribution for single rare variant tests

To generate an empirical null distribution for single rare variant tests, phenotype permutation was performed using PLINK, where case/control labels were randomly re-assigned in each permutation. By using the PLINK command --mperm-save-all, we saved all permuted test statistics. These statistics were combined together into a single vector, ordered, and split into bins where the size of each bin was the number of permutations. To average across permutations, the median *p*-value was pulled from each bin. We used 100 permutations to derive the empirical null.

### Type I Error and Power for gene-based tests

We assessed type I error and power for gene-based tests using the following 5 models: 1) SKAT with empirical significance, 2) SKAT with random permutation for significance, 3) SKAT adjusting for PCs of ancestry as covariates, 4) SKAT adjusting for PCs of ancestry as covariates with random permutation for significance, and 5) SKAT modified with Bootstrat. To simulate the effect of population structure, we randomly selected 70% of American samples and 30% of Swedish samples as cases and set the remaining samples to be controls. Type I error was assessed at the 0.05 with 1,000 iterations and at lower significance levels (0.01, 0.005, 0.001 and 0.0005) with 20,000 iterations under the null models (odds ratio of 1). Power was assessed at the 0.05 and 0.01 significance levels with 1,000 iterations for each model. We assigned each variant to be neutral or have an effect with a 30% (or 50%) probability and simulated the phenotype based on the genotypes using odds ratios of 2.5 and 5. We randomly sampled genes on chromosome 1 for both type I error and power assessments.

## Acknowledgments

GMP is supported by the National Heart, Lung, And Blood Institute of the National Institutes of Health under Award Number K01HL125751. The content is solely the responsibility of the authors and does not necessarily represent the official views of the National Institutes of Health. BMN acknowledges funding from the Gerstner family foundation. BR and KB acknowledge funding from the Max Planck Society. Carl-Johann SIMON-GABRIEL is supported by a Google European Doctoral Fellowship in Causal Inference. The funders had no role in study design, data collection and analysis, decision to publish, or preparation of the manuscript.

## Supporting Information

**S1 Fig. Selection of the principal components.** QQ plot of the test statistics of common variants when various numbers of principal components were used.

**S2 Fig. Selection of the Bootstrat parameter.** QQ plot of the test statistics of common variants for *γ* ranges between -3 and 8.

**S1 File. Psuedocode for estimating the Bootstrat parameter.**

